# Brain rewiring during developmental transitions: A Comparative Analysis of Larva and Adult *Drosophila melanogaster*

**DOI:** 10.1101/2024.05.01.592061

**Authors:** Prateek Yadav, Pramod Shinde, Aradhana Singh

## Abstract

The brain’s ability to adapt through structural rewiring during developmental transitions is a fundamental aspect of neuroscience. Our study conducts a detailed comparison of *Drosophila melanogaster* ‘s brain networks during larval and adult stages, revealing significant changes in neuronal wiring during developmental phases. The degree distribution of the larval brain deviates significantly from power-law behavior and fits well with the Weibull distribution. In contrast, the adult brain exhibits power-law behavior in its degree distribution, with the exponent for the out-degree distribution lying in the scale-free regime and the exponent for the in-degree distribution being close to this regime. This difference reflects a change in the robustness of brain development from larval to adult phases. The core of these networks also changes during development in terms of their cell composition and topological influence. The larval core comprises Mushroom Body neurons, while the adult core mainly has Antennal Lobe neurons. Moreover, all the core neurons in the larval brain are also part of the rich-club neurons, a group of neurons with high in/out degrees that are well connected, whereas the same is not true for the adult brain network. Additionally, the core of the larval brain displays a more heterogeneous connectivity profile in its second-order neighbors compared to adult brain neurons, indicating greater diversity in larval brain connectivity. Our work stands as a step forward in understanding the rewiring of brain networks across the life stages of *Drosophila melanogaster*.

## 1 Introduction

Connectomics is the comprehensive study of neural connectivity patterns, encompassing both structural and functional relationships across various spatial scales within the nervous system [1], [2]. As the brain performs tasks through the coordination between the different neuronal cells, studying its rewiring is useful in understanding how neurological structures translate to function [3]. The mapping of the brain assigning the links between the different components of the brain including neurons, cells, or different brain regions through axons, fiber tracts, or temporally correlated activation [4] mainly relies on experimental techniques such as diffusion-weighted magnetic resonance imaging (DW-MRI), functional magnetic resonance imaging (fMRI), electroencephalography (EEG) for macroscale studies, serial sectioning of nervous tissue, and electron microscopy (EM) for microscale studies [5]. To date, comprehensive brain mapping that reveals connectivity patterns at the cellular level has been achieved for only a few species, one of which is *Drosophila melanogaster*, commonly known as the fruit fly. Drosophila is an exemplary model organism for connectomics research due to its extensive use in genetics and molecular biology. The substantial knowledge and comprehensive experimental protocols associated with *Drosophila melanogaster* biology make it a valuable resource for connectomics investigations. Additionally, *Drosophila* exhibits a diverse repertoire of well-characterized behaviors, further enhancing its suitability as a model organism for studying the neural circuitry underlying behavior and cognition [6], [7].

The complete metamorphosis of higher insects including *Drosophila* is not only fascinating but also shrouded in mysteries, particularly those related to the nervous system. The smooth transition from the larval stage to a morphologically and physiologically distinct adult form in *Drosophila* necessitates complex alterations in the nervous system [8], [9]. Such as the neuronal reorganization in the development of *Drosophila* from the larval to the adult phase preserves some cell types, such as Kenyon cells and antennal lobe neurons, which are involved in learning, memory, and olfaction and are found in both stages. However, certain types of neurons that are associated with metamorphosis, mating, hormonal regulation, etc., are only found in the adult brain [10–12]. Also, the moonwalker descending neuron (MDN) circuit and associated Pair1 interneurons responsible for locomotion in the larval stage, undergo remodeling during metamorphosis [13]. With the generation of the complete connectomes of the larval and adult forms of the fruit fly [10, 11], we got an opportunity to compare the two brain networks and to see what insights network science offers when applied to these entire datasets in their comparison.

Network science is a powerful tool for unveiling the hidden characteristics of complex systems in terms of structural and functional relationships between their constituents using a graph representation [14]. Herein, neurons are defined as nodes, and synapses between neurons are defined as edges. Network framework provides a cue into whether the structural environment confers opportunities for or constraints on individual node action. The degree of a node, which is the total count of its neighbors, determines its centrality and is therefore the basic metric to characterize a given network. A node with a high degree is more influential. Moreover, cohesive groups of influential nodes have been discovered in various brain networks in the form of network-core and rich-club organization [15, 16]. The k-core decomposition constitutes a process of separating the nodes in a graph into groups based on the degree sequence and topology of the graph [15],[17]. Core decomposition analyses have revealed interesting insights in the field of neuroscience. A k-core decomposition analysis carried out separately for in and out-degree revealed that the k-core for in-links consisted predominantly of motor neurons while the k-core of out-links consisted comprised primarily of sensory neurons in the brain of *C. elegans* [18]. A study of Human functional brain networks using fMRI revealed that the k-core comprises the brain regions that are active in the subliminal state of the brain [19]. Similarly, rich clubs have been reported in the connectomes of *C. elegans* [20], cat [21], [22] and mouse [16]. In human brain networks, a rich club comprising cortical regions (superior parietal, precuneus, superior frontal cortex) as well as sub-cortical regions (putamen, thalamus, and hippocampus) from both hemispheres was found [23]. The involvement of disrupted rich club connections has been linked to various neurological and pathological disorders, providing insights into their underlying pathologies [24],[25],[26]. Both k-core decomposition and rich club analyses offer a mesoscopic view of how influential nodes interact with each other and impact the rest of the network.

Our study aims to develop a network-based framework to enhance the system-level understanding of the nervous system metamorphosis in *Drosophila*. Particularly, we aimed to compare and contrast neuronal connectivity between larval and adult *Drosophila* brain networks based on their degree distribution and the core decomposition. We study the D-core, which is the directed counterpart of the k-core decomposition [27]. For this, we study two publicly available datasets: one that offers the whole brain connectome of a female fruit fly larva (first instar) [10] and another that encompasses the entire brain of an adult female fruit fly (Flywire dataset v630) [28, 29]. We find some similarities between the two networks, such as both are modular and are not strongly connected, also, some differences, such as the larval brain is weakly connected but the adult brain is not. Also, we find that the degree distribution of the larval brain is stretched exponentially and fits well with the Weibull Distribution [14], whereas the adult brain exhibits a strong adherence to a power-law distribution in both the out-and in-degree distributions, with their exponent being in and very close to the scalefree regime respectively. Furthermore, we find that the core of the brain network remodels during metamorphosis, as follows: (a) The core of the larval brain consists of mushroom body neurons, which are involved in learning and memory. In contrast, the core of the adult brain comprises antennal lobe neurons, which process olfactory information. (b) The larval brain displays a strong rich-club organization comprising the high-degree nodes, which are also the core neurons, the same is absent in the adult brain. (c) The larval brain network exhibits a more globally influential core, likely due to the integrative multi-sensory computations performed by the constituent mushroom body (MB) neurons. In contrast, the core of the adult brain is more localized in terms of its overall contribution to the clustering and efficiency of the whole network. This analysis provides a direction to understand and capture important developmental changes in an organism’s brain.

## 2 Results

### 2.1 Distinct structural properties across *Drosophila* developmental stages

The structural properties of the adult and larval *Drosophila* developmental stages were assessed, and we noted peculiar differences between these two brain networks (see Table 1). At first glance, the adult and the larval brain networks differed in their basic network properties. The adult brain is much larger than the larval brain, with over 40 times the number of nodes (neurons, *N*, see Table 1). This difference in size could be attributed to the adult brain’s need to support much more complex behaviors such as foraging, navigation, and mating (cite). The larval brain network exhibits a much higher connection density (*d*) and clustering than the adult brain, which makes it weakly connected (Table 1), the same is not true for the adult brain. However, the adult brain has a higher modularity (*Q*, Table 1) than the larval brain, suggesting that the adult brain network has densely connected components performing more independent specialized functions as compared to the larval network. Moreover, these basic network properties also reflect similarities between these two brain networks. Both the larval and adult brain networks exhibited smaller average clustering coefficients, indicating that these networks are not strongly connected. This is reflected in the results of the Strongly Connected (SC) test, which were negative, as shown in (Table 1). A directed network is strongly connected if there is a path from every node to every other node. Also, both networks have a higher value of modularity (*Q*), suggesting a strong modular structure with well-partitioned communities that are densely connected within themselves but less connected. We further explored the network structures of both networks in detail.

**Table 1:** Network properties of Adult and Larva Connectome. Here, *N, N*_*c*_, *d*, ⟨*K*⟩, ⟨*C*⟩, *Q*, SC, and WC represent the number of nodes, the number of edges, connection density, the average degree, the average clustering coefficient, modularity of Louvain partitions with the resolution parameter 1, strongly connected components, and weakly connected components, respectively.

| Network | $N$ | $N_c$ | $d$ | $\langle K \rangle$ | $\langle C \rangle$ | $Q$ | SC | WC |
| --- | --- | --- | --- | --- | --- | --- | --- | --- |
| Adult | 124891 | 2613129 | 0.0002 | 42 | 0.11 | 0.66 | False | False |
| Larva | 2952 | 110140 | 0.01 | 75 | 0.18 | 0.52 | False | True |

To gain deeper insights into the organization of connections within brain networks, we further analyze the degree distribution of these networks. We found an intriguing observation regarding the degree distribution for both in-degree and out-degree in adult and larval brain networks. The degree distribution exhibits two distinct behaviors in the initial and tail segments. In the larval degree distribution, the initial segment displays a noticeable plateau, representing a significant proportion of the overall distribution. In contrast, such a distinction is less pronounced in the adult network (Fig. 1). This observation suggests unique structural characteristics in the larval stage compared to the adult stage. The presence of a fat tail in the degree distribution of both networks indicates the existence of hubs. Previous studies have demonstrated that many real-world networks, such as the frequency distribution of earthquake magnitudes, word frequency, citations, telephone calls, the intensity of wars, and the net worth of wealthy Americans, exhibit hub-like structures, as indicated by the flat tail in their degree distributions [30]. These flat tail distributions often exhibit a power-law behavior, where the probability function for the degree distribution is given by the form *P* (*k*) ∝ *k*^−*γ*^, where, *P* (*k*) refers to the probability of finding a node with degree *k* in the network and *γ* is the power law exponent. For the exponent 2 *< γ <* 3, the scale-free property emerges as the second moment of the degree distribution diverges [30]. Consequently, we decided to examine whether the degree distributions of fly connectomes fit a power law.

**Figure 1:**
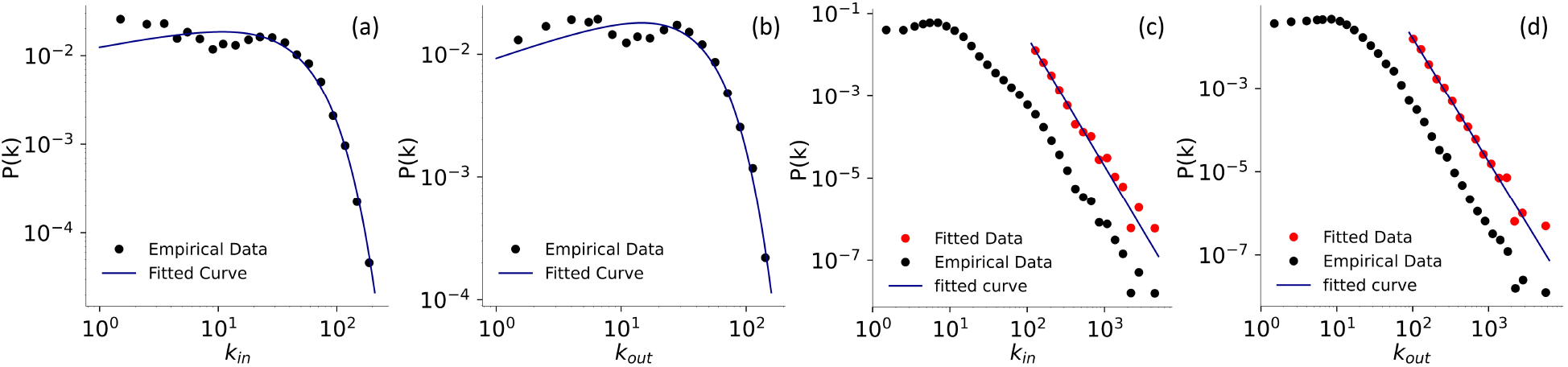
The in-degree and out-degree distribution of the larval and adult brain networks on a log-log scale. Subplots (a) and (b) show the in-degree (*k*_in_) and out-degree (*k*_out_) distributions of the Larva Connectome, while subplots (c) and (d) show the corresponding distributions for the adult brain network. In subplots (a) and (b) for the larval brain, the distributions are fitted with the Weibull distribution, while in subplots (c) and (d) for the adult brain, they are fitted with the power-law distribution using the maximum likelihood method mentioned in the method section. In subplots (c) and (d), the black points represent the probability distribution function for the empirical data, while the red points represent the probability distribution function of the fitted data (which includes degree values ≥ *x*_min_). Further details on the minimum degree fitting with the power law are provided in Table 1 of the supplementary material.

We observed that for the larval brain network, the degree distribution is stretched exponentially, and the proportion of data following the power-law fit is very small (Fig. 1 (a), (b) of Supplementary material). Both in-and out-degree fits well with the Weibull distribution (Fig. 1 (a), (b)). Weibull is a mixture of the exponential and the power-law, and therefore the probability function for the Weibull distribution has two parameters and is given as 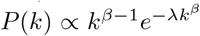 [31]. For the parameter *β* = 1, it becomes pure exponential. We use maximum likelihood estimation, as mentioned in the method section, for this fitting. The entire range of the in/out-degree of the larval brain network fits well with the Weibull distribution, confirmed by the low Kolmogorov-Smirnov distance values 0.0396 and 0.0388, respectively. The distribution parameters *λ* and *β* are determined to be 0.0247 and 1.2409 for in-degree and 0.0244 and 1.3348 for the out-degree. Weibull has also been found to be a better fit for the degree distribution of the functional human brain network [32]. The parameter *β >* 1 for both the in-and-out-degree distributions indicates that the failure rate of this fit increases as the network grows. This indication matches with our observation that as the fly evolves in time and becomes a bigger, fully mature adult, and the brain network grows, both the in- and out-degree distribution fits poorly with the Weibull distribution (Fig. 1 (c), (d) of Supplementary material). Instead, both the in-and out-degree of the adult brain network fits well with the power-law distribution, with most of the data falling on the straight line fitting the distribution on the log-log plot (Fig. 1 (c), (d)). Detailed statistics of this fitting can be found in Table 1 in the supplementary material. It is noticeable that the exponent of the power-law for the out-degree of the adult brain is in the scale-free regime (*α* = 2.96), and for the in-degree, it is very close to the scale-free regime (*α* = 3.15), indicating that the existence of scalefreeness in this network which is missing in the larval network. This suggests that as the Fruit fly brain grows, scalefreeness emerges to provide better resilience to the errors in the network Moreover, adherence to a power law in the adult brain networks indicates a higher prevalence of putative hubs. In contrast, in the larval network, only high-degree neurons follow the power law, suggesting the presence of smaller quantities of the putative hubs compared to the adult network. Such patterns likely reflect underlying evolutionary processes that influence network formation and stabilization in the adult brain.

To delve deeper into these network differences, we employ rich club and D-core analyses, which provide enhanced insights into the intricate organization of connections among high-degree neurons and the core structures within each network, highlighting their hierarchical and functional organization.

### 2.2 D-core decomposition and rich club analyses

Using the method described in the method section, we obtained the set of nodes belonging to the Frontier D-cores for both the larval and adult connectomes. Despite the adult brain is much larger than the larval brain, it loses connections more rapidly during the core decomposition process due to its lower connection density compared to the larval brain network. As depicted in Fig. 2 (b, c), in the D-core (10, 8), the adult brain loses almost 80% of its neurons, whereas, in the same D-core, the larval brain loses approximately 20% of its neurons. Therefore, despite the adult brain network having more high-degree neurons than the larval brain, the number of nodes in their Frontier D-core is similar. Additionally, the heterogeneity of cell types in the Frontier D-core is similar for both brain networks (Fig. 2 of the Supplementary material). These Frontier D-cores being the innermost D-cores with nonzero cells and having the high-connectivity neurons serve as the core of the network. From now on, we refer to these as the core of the network. To gain a comparative understanding of the core of the larval and the adult brain networks, we examine the compositions of the core neurons and their neighbors in detail in the following section.

**Figure 2:**
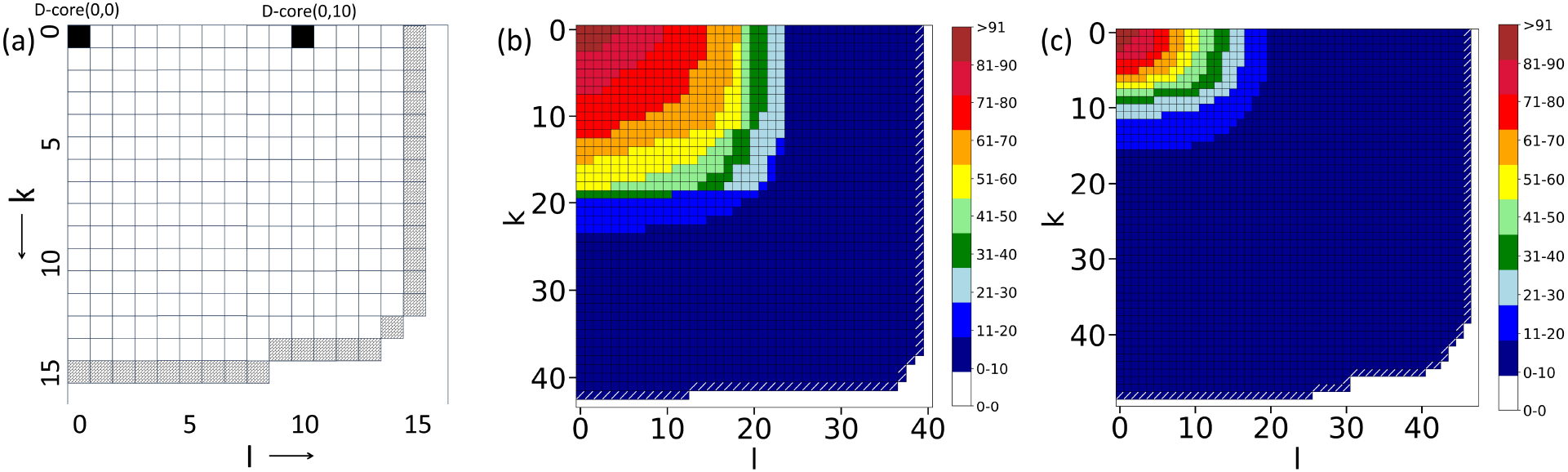
D-core decomposition of the networks: (a) A schematic of the D-core matrix with the frontier cells filled with diagonal brick patterns. Each (k,l) cell represents a D-core(k,l). Here, *k*_max_ and *l*_max_ are both 15. (c) D-core matrix of the Adult Fly Connectome. (b), (c) plots the D-core matrix of the Larval and Adult Fruit-fly Connectomes respectively. In (b) and (c), frontier cells are highlighted by white-colored oblique lines.

#### 2.2.1 The larval core is comprised of mushroom body neurons

In the larval brain, the core consists of 181 neurons, including 114 Kenyon Cells (KCs), 44 Mushroom Body Output Neurons (MBONs), 21 Mushroom Body Input Neurons (MBINs), and 2 Local Interneurons of the Anterior Paired Lateral Cell (Figure 3 (a)). All of these neurons belong to the mushroom body, which has been conventionally perceived to primarily be a center for olfactory memory and has also been implicated in myriad other behavioral tasks, namely odor preference, taste preference, and associative learning between odor and taste [33, 34]. Each of the neuron types found in the core has a particular role in the functioning of the memory circuit. KCs, located in the calyx of the mushroom body, obtain Olfactory sensory information from the sense organs [35, 36] and send outputs to other KCs, MBINs, and MBONs [36, 37]. On the other hand, the gustatory information from the same chemosensory apparatus that includes the olfactory sensory system is relayed to the MBINs [35, 36]. MBINs establish synaptic connections at both the presynaptic and postsynaptic terminals of multiple KC-to-MBON synapses [37]. They are composed of DANs, Octopaminergic neurons, and a few neurons whose neurotransmitters haven’t been identified yet [36]. MBONs are output neurons that are responsible for generating motor output based on the olfactory and gustatory information [36]. They receive input from KCs, MBINs, other MBONs, and non-MB neurons [37].

**Figure 3:**
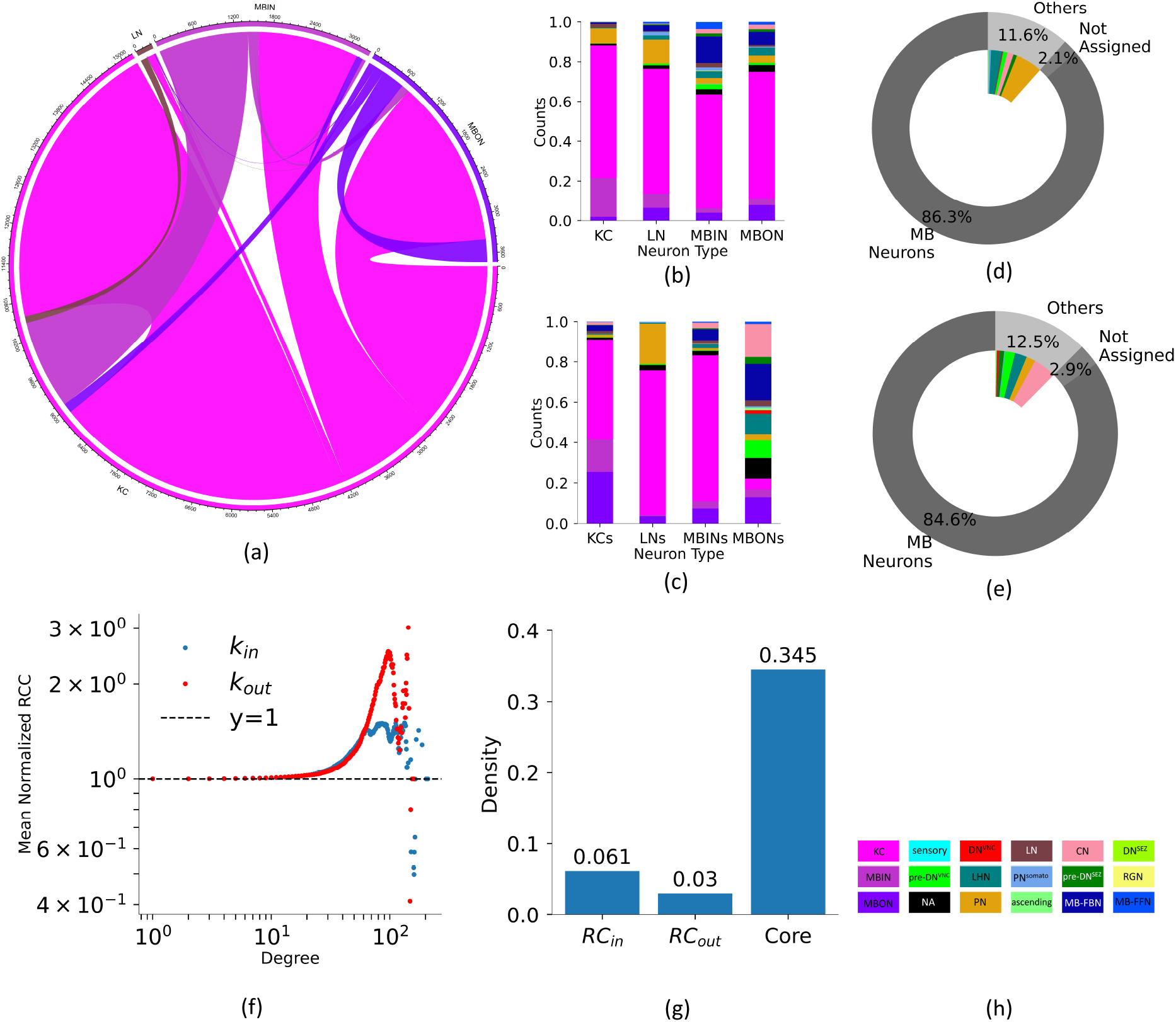
Analysis of the core of the larva connectome. The chord diagram in (a) plots the neuron types present in the core and the connections among them. In (b) and (d), the normalized cell type composition of predecessors and successors of the core are plotted. Pie charts in (c) and (e) summarize the type composition of predecessors and successors of the core neurons. (f) plots mean normalized rich club coefficient vs. degree, bars in (g) plot connection density of the in-degree RC, out-degree RC, and core, and (h) presents the color code legend for the different cell types present in predecessors and successors of the core neurons.

Furthermore, we explored which neuronal cell types are directly connected with the core neurons, as these neurons act as a bridge between the other neurons and the core. Neurons that are postsynaptic to frontier neurons are termed their successors, while presynaptic to frontier neurons are termed their predecessors. All four types of core neurons are mostly connected to the KC neurons, followed by MBIN neurons, and then by MBON neurons, which are the MB neurons (Fig. 3 (b)). Additionally, there are connections to MB-FBNs, and MB-FFNs, all of which belong to the same neuropil as the frontier neurons, thus demonstrating homogeneous connectivity. In total, these MB neurons account for 86.3% of the neighbors of the core, indicating that the nearest neighbors of core neurons are similar types of neurons. The inter-specific types include Projection Neurons, CNs, pre-DNs^VNC^, and pre-DNs^SEZ^. This information is well summarized by the pie chart in Fig. 3 (d). The successors of the core also mostly exhibit a similar type of composition pattern as shown by the predecessors. However, the successors of the core differ from predecessors in having a greater type diversity among the neighbors of the core MBON neurons (Figure 3 (c), (e)), indicating that these connections likely play a role in generating behavioral responses.

Furthermore, we performed a rich club analysis to gain deeper insight into the core of the larval brain.

This analysis focuses on a group of high-degree nodes that are more interconnected than their corresponding random surrogates. We calculated the rich club coefficient (*RCC*) as defined in the method section for the corresponding 100 degree preserved networks and the normalized *RCC* obtained for the larval brain network by this. This normalized *RCC* is plotted in Fig. 3 (f). We find that the larval connectome shows prominent rich club organization for both in and out-degree (Fig. 3 (f)). The density of the in-degree rich club neurons and out-degree rich club neurons is 0.061 and 0.03, respectively, which is 6 times and 3 times, respectively, the whole network’s density (Fig. 3 (g)). Further analysis indicates that all neurons within the D-core are also part of the rich-club. The density of these D-core neurons, at 0.345, is approximately 30 times greater than that of the entire network. This significant increase in connectivity within the D-core compared to other rich-club neurons suggests that the D-core represents a more intensely connected subgroup. Essentially, it could be considered the “rich-club of the rich-club,” highlighting its critical role in network communication and function (Fig. 3 (g)).

#### 2.2.2 The core of the Adult fly brain is comprised of Antennal Lobe (AL) neurons

In the adult brain, the frontier D-cores comprise 198 neurons. Of these, 122 are Local Interneurons, 74 are Projection Neurons, and 2 are Antennal Lobe Input (ALI) neurons (Figure 4(a)). All these neurons belong to the antennal lobe, which serves as the primary center for olfaction in the fly brain. The antennal lobe of the fly is analogous to the olfactory bulb of vertebrates. Here, olfactory information from the antennae and maxillary palp reaches the Antennal Lobe Local Interneurons (ALLNs) and Antennal Lobe Projection Neurons (ALPNs) [35]. After processing in the antennal lobe, the information is relayed to the neurons in the Mushroom Body and lateral horn (LH) [35]. A little is known about ALI neurons, these neurons are labeled as AL-MBDL1 neurons in the hemibrain connectome dataset [38], and they are thought to be involved in sending feedback signals to the antennal lobe. Additionally, they are likely influenced by the activity of Mushroom Body and lateral horn neurons, as they exhibit arborizations in the ring neuropil, which is connected to these two regions [39]. The exclusive presence of antennal lobe neurons in the frontier D-cores could be because of the pivotal role of olfaction in the ecology of an adult fruit fly—locating food sources, finding oviposition sites, avoiding predators, and selecting mates.

**Figure 4:**
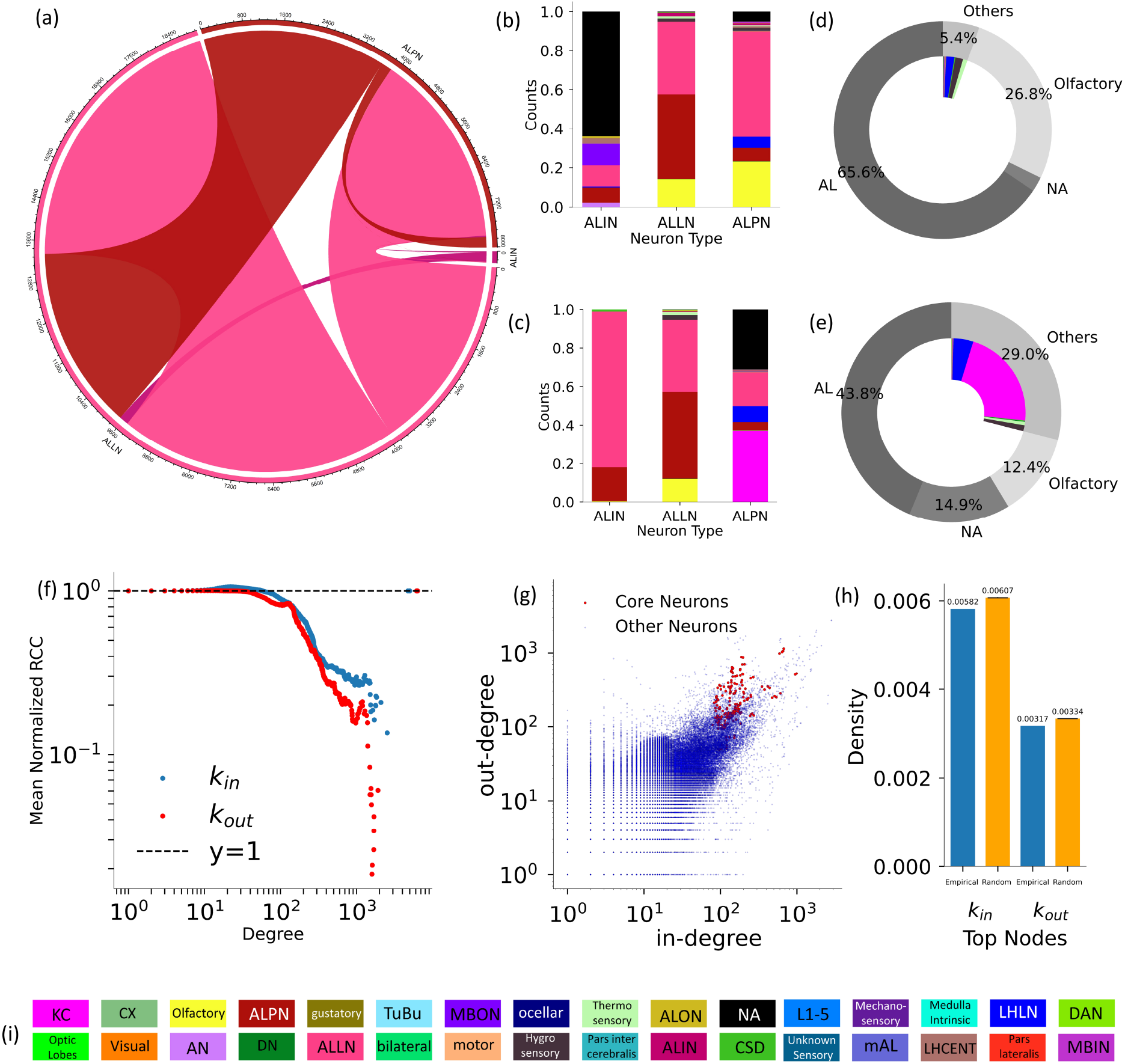
Analysis of the core of the Adult brain connectome. The chord diagram in (a) illustrates the neuron types present in the core and the connections among them. Figures (b) and (d) display the type composition of predecessors and successors of core neurons, respectively, while the corresponding pie charts are summarized in (c) and (e). Figure (f) plots the mean normalized rich club coefficient versus degree. Figure (g) illustrates the in-degree versus out-degree for all neurons, highlighting the values for the core neurons. Additionally, Figure (h) depicts the density of the top 4.9% high in-degree neurons, top 8.5% out-degree neurons, and the corresponding degree-preserved randomized networks. Finally, Figure (i) provides the color code legend for cell types observed in (b) and (d).

To understand the functional role of the frontier neurons, we investigated the neuronal types that interact with them. Excluding unassigned neurons, most of the in-connections are from other Antennal Lobe Local Interneurons (ALLNs) and Antennal Lobe Projection Neurons (ALLPNs) for all these three types of core neurons (Figure 4 (b)). Thus, all three types show homogeneity in their in-connections, as their predecessors belong to the same neuropil, the Antennal Lobe. We observe that the interspecific connections primarily involve Lateral Horn Local Neurons (LHLNs) and olfactory neurons, with a smaller proportion connecting to Mushroom Body Output Neurons (MBONs). This is due to the limited number of Antennal Lobe Interneurons, which also explains why unassigned types occupy a small region in the pie chart (Figure 4 (d)). The pie chart further illustrates this, showing a higher contribution from hygrosensory and thermosensory neurons than MBONs among the inter-specific connections. Notably, in the pie chart, the olfactory type is shown separately because it is composed of Olfactory Receptor Neurons (ORNs), which have a close association with the Antennal Lobe. After their odorant receptors bind to odor molecules, ORNs send signals to the local neurons and projection neurons of the antennal lobe [40, 41]. Regarding the out-connections, we see a similar dominance of ALLNs and ALPNs albeit a little diminished because of the presence of Kenyon cells as out-neighbors of the ALPN type frontier neurons (Figure 4 (c)). This is made clear when comparing the two pie charts (Figure 4 (d) and Figure 4 (e)). This is because PNs deliver their sensory input to the Kenyon cells (KCs) of the Mushroom Body calyx [42, 43]. After the KCs, just like in the in-connections, we see a presence of LHLN neurons.

Further, we found that the core of the adult brain consists of the top 4.0% and 8.5% of high in- and out-degree nodes, respectively. These core nodes do not form a rich club, as the normalized RCC takes a value lower than 1 for higher degree values Fig 4(f). The in-and-out-degree of the core nodes are highlighted in Fig 4(g). However, we find that the normalized *RCC* takes a value greater than one at the intermediate values of the degree, as also shown in the article [11]. This article [11] considers rich-club neurons as all neurons with degrees above a specified intermediate value where normalized RCC takes a value greater than 1. Using this criterion, we identified a substantial Rich Club comprising 26, 662 neurons. However, it is important to note that, according to the definition of a rich club, simply having a high degree does not necessarily qualify nodes as members of the rich club. Here, the indication of the normalized RCC having values lower than 1 indicates the absence of this phenomenon for the high-degree nodes of which the core is part. This is also confirmed by the lower density of the connections among the neurons that are in the top 4.9% and 8.5% of high in- and out-degree nodes, respectively than their degree preserved random surrogates (Fig.4 (h)). In summary, the pronounced rich club organization in the larval network and its absence in the adult network align with their respective average clustering coefficient values. The larva’s higher average clustering coefficient ⟨*C*⟩, as referenced in Table 1, indicates a greater tendency to form tightly connected, cohesive groups. This observation underscores the developmental and structural differences between the larval and adult stages.

### 2.3 Type compositions of ‘neighbors-of-neighbors’ of Frontier D-cores

The study thus far has demonstrated that both larval and adult core neurons primarily exhibit a similar pattern, characterized by a high proportion of intra-specific connections. Building on these findings, we now aim to explore further by comparing the type diversity of their second-order connections, specifically examining the neighbors of their neighbors to gain deeper insights into the network’s structure. We find that 25% of the second-order neighbors of the larval core are MB neurons, while 10% of the second-order neighbors of the adult core are AL neurons. At this level, the cell composition differs significantly between the larval and adult brains. The larval connectome exhibits greater type diversity compared to the adult connectome among both predecessors and successors of immediate neighbors of the core (Fig. 5).

**Figure 5:**
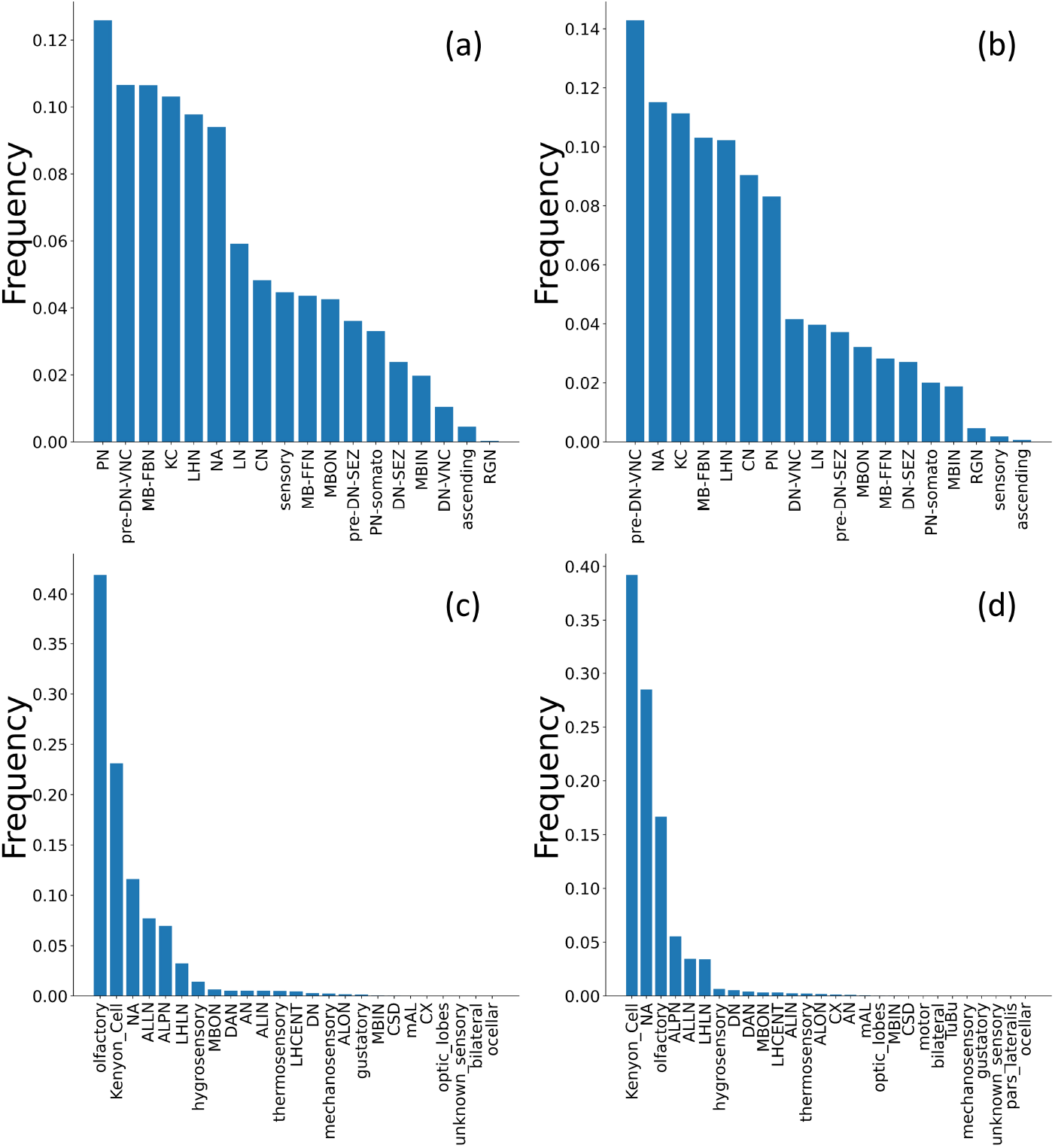
Normalized Frequency of Different Cell Types in Second-Order Neighbors of the Core. Panels (a) and (b) depict the normalized frequency of predecessors and successors of immediate neighbors of the larval brain core, respectively. Likewise, panels (c) and (d) illustrate the normalized frequency of predecessors and successors of immediate neighbors of the adult brain core, respectively.

To further quantify this diversity, we calculated both Shannon entropy and Simpson’s index (see Methods section) to measure the diversity in distribution by considering both type richness and abundance. As shown in Table 2, the higher values of both indices for the larval connectome indicate greater diversity in terms of its second-order neighbors.

**Table 2:** Measuring the diversity of the second-order connections. The larval connectome exhibits greater diversity in second-order connections compared to adult connectomes, as evidenced by higher normalized Simpson and Shannon indices.

| Network | Connection type | Simpson index | Shannon index |
| --- | --- | --- | --- |
| Adult | Predecessors of predecessors | 0.16 | 0.13 |
|  | Successors of successors | 0.13 | 0.11 |
| Larva | Predecessors of predecessors | 0.67 | 0.22 |
|  | Successors of successors | 0.62 | 0.21 |

The adult brain shows high second-order associations with olfactory neurons and Kenyon cells, indicating the central role of core neurons in olfaction. In contrast, for the larval brain, the second-order associations are primarily to the pre-descending ventral nerve cord (VNC) neurons, projection neurons (PN), MB feedback neurons (MB FBNs), mushroom body output neurons (MBONs), lateral horn neurons (LNs), and convergence neurons (CNs). The pre-descending ventral nerve cord (VNC) neurons play a vital role in integrating and transmitting nerve signals, coordinating sensory input and motor output, and controlling muscle activities in the periphery [44]. Projection neurons transmit diverse sensory modalities, including olfactory, mechanical, thermal, and gustatory information [10]. MB feedback neurons (MB FBNs) provide feedback from MBONs to help integrate aversive and appetitive memory apparatus [45]. Lateral horn neurons (LHs) are involved in coding odor valence and contributing to innate odor responses [46]. Convergence neurons (CNs) integrate inputs from both the mushroom body (MB), representing learned values and the lateral horn (LH), representing innate values [10, 46].

This disparity may reflect the broader functional roles played by mushroom body neurons in the larval stage compared to the more focused role of the antennal lobe in processing chemosensory inputs in the adult stage.

Based on the diversity of the types of neighbors’ neighbors of core neurons, it can be inferred that the larval core is more generalized, connecting with a broader array of diverse cell types. In contrast, the adult frontier core appears to be specialized, with connections extending to a more limited range of cell types. This suggests that the larval network may be more adaptable, while the adult network is optimized for specific functions.

### 2.4 Topological roles of the Frontier D-cores neurons

To further investigate the role of core neurons in the larval and adult brain networks, we examine the impact of their removal on the average clustering coefficient of the entire graph and their role in providing the shortest path between nodes of the largest strongly connected component. This is achieved by conducting targeted and random attacks on both brain networks and observing changes in clustering, and global efficiency.

In a targeted attack, all frontier neurons are eliminated simultaneously, while in a random attack, an equivalent number of nodes to those in the set of frontier neurons are randomly selected and eliminated at once. The clustering parameters are evaluated using 1000 random attacks for both the larval and adult networks. However, for efficiency, 1000 random attacks are exclusively applied to the larval network, while for the adult network, only 10 attacks are used due to the computational complexity of the directed efficiency parameter and the larger size of the adult network, requiring a reduced number of iterations for practical feasibility.

The bar plots in Figs. 6(a) and (b) compare the ratio between the clustering and global efficiency of the network before and after random and targeted removal of nodes in the larval and adult brains, respectively. It shows a significant difference between the effect of random and targeted attacks on the clustering coefficient and efficiency of the larval brain. However, no such difference is observed in the adult network. Moreover, the mean normalized betweenness centrality of the core nodes in the larval brain is 8 times higher in the larval brain than in the adult brain (Fig. 6 (d)), indicating a greater contribution of the core neurons in the larval brain to its overall efficiency than in the adult brain and hence having a central role in the network.

**Figure 6:**
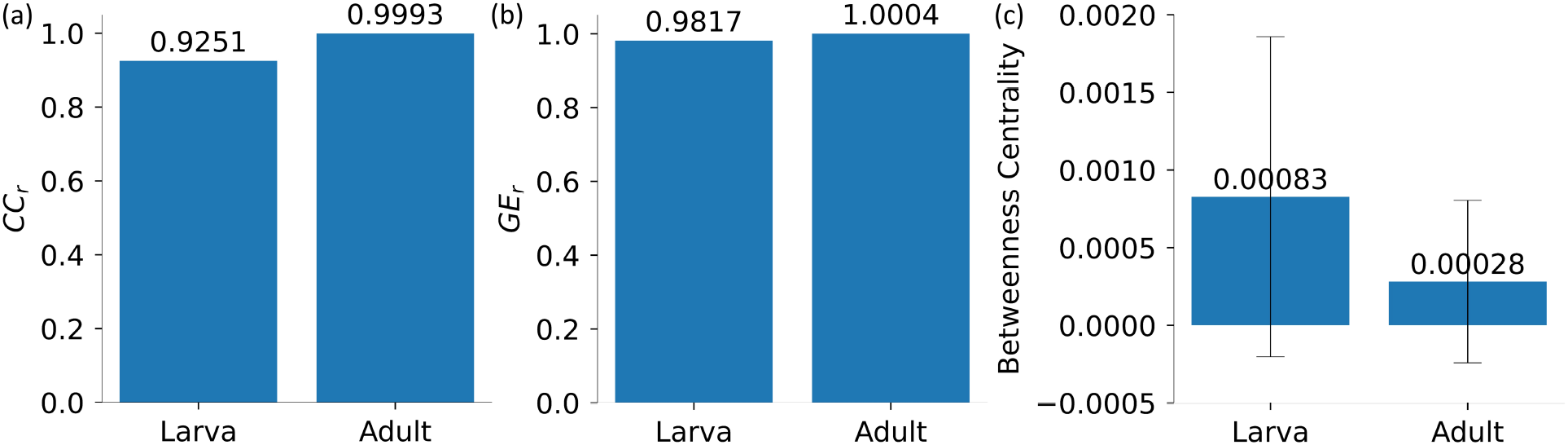
Topological role of the frontier D-core neurons in Larval and Adult brain. Plots (a) the ratio of the clustering, (b) the ratio of the efficiency of the brain networks, before and after removal of the D-core neurons and (c) plots betweenness centrality of the Frontier D-cores neurons of the Larval and the Adult brains. The statistical significance is done using permutation testing involving 100, 000 permutations. The error bars are shown in black.

This comparatively greater topological significance of the core neurons in the larval network could be because, in the adult, the cores are more limited in their functional scope, as revealed by the type composition of the neighbors of their neighbors in Fig. 5.

## 3 Discussion

We compared the larval and adult fruit fly neuronal brain networks to analyze the developmental changes occurring during the growth of the fly brain throughout metamorphosis. The process of brain rewiring during fly development involves profound metamorphosis characterized by neuronal death, brain reorganization, and the development of adult-specific neurons. Therefore, larval and adult life stages are expected to differ significantly, each linked to stage-related functions and environmental stimuli.

Our study aims to pinpoint the major changes that occur during the developmental process. We find significant alterations in the mesoscopic organization of brain connectivity as *D. melanogaster* evolves from the larval to the adult phase. However, we also notice universal similarities between the two brain networks, both networks exhibit sparsity and a modular organization likely preserved through evolutionary processes, which were also previously reported [47]. We find that the adult brain is more modular than the larval brain, reflecting an enhancement of specialized brain compartments with distinct functions. This enhancement is also reflected in the reduction in density, clustering, and the disappearance of weakly connected properties observed in the larval brain during the adult phase. Overall, this suggests that the adult brain has evolved to feature specialized structural compartments, which likely serve as functional epicenters. This organized integration has been previously reported and demonstrates how brains can systematically process sensory information into structured feature maps [48]. These maps form diverse pathways that act as essential links between vision and behavioral controls, highlighting the brain’s ability to coordinate complex functions efficiently.

Moreover, the adult degree distribution displays a stronger conformity to a power-law, with its out-degree distribution falling within the scale-free domain and the in-degree distribution narrowly failing to meet the threshold. In contrast, the larval degree distribution fits with the Weibull distribution, with its growth parameter (*β*) being greater than one. This indicates that as the network grows, it adapts to have the scale-free property to gain robustness against failures. A recent study of the brain network of *C. elegans* across its development found that degree distributions consistently did not follow a power-law model across all stages [49]. This observation underscores the intricacies inherent in the developmental process of *Drosophila* compared to *C. elegans*, a distinction that aligns with the fact that the former undergoes complete metamorphosis.

Further, analyzing the core of larval and adult brain networks using D-core decomposition, which is analogous to the k-core of undirected networks, we find that the larval core comprises Mushroom Body neurons, while the adult core has Antennal Lobe neurons. The different constituents of the core for the adult and larval connectome could indicate their disparate lifestyles. The exclusive participation of Mushroom Body (MB) neurons in the larval brain core indicates that the MB neurons have a central role in its functioning. This is also supported by the recent findings [35, 37, 50] finding the Mushroom body is not only the center of olfactory learning but also in integrative multi-sensory learning and memory, given the fact that the Kenyon cells receive thermal, gustatory and visual input through non-olfactory Projection neurons and that such multi-sensory Kenyon cells and MBINs synapse onto the MBONs.

Our analysis of the neighbors of the core provides an understanding of the neuronal pathways and the role of different types of neuronal cells in forming a complex yet robust network architecture. We find that the cell type composition of the core is preserved in the immediate neighbors in both the larva and adult as the core neurons and their immediate neighbors belong to a single neuropil: Mushroom Body in the former and the Antennal Lobe in the latter. Moreover, the larval network displayed a strong rich club organization for both in and out-degree. The core being part of the rich club is more connected than in general all other rich club neurons, indicating the existence of a “rich club within the rich club.” This indicates the hierarchical organization of connectivity, which may have implications for functional integration and information processing.

To further analyze the connectivity patterns of the two frontier cores, we examine their second-order connections, namely the neighbors of neighbors. This analysis reveals a higher type diversity in the connections of the larval connectome for both predecessors and successors, indicating that the core of the larval connectome has a global influence on the network. This is supported by a significant decrease in the average clustering coefficient and global efficiency of the larval connectome when we remove the core nodes. Conversely, the neighbors of the neighbor of the adult core are predominantly involved in olfaction. Consequently, the adult core, specialized in the olfactory domain, exerts a peripheral influence, as evidenced by the negligible effect on its average clustering coefficient and global efficiency after the loss of core nodes, as well as a lower mean betweenness centrality of its core neurons.

The larval connectome’s globally influential core may be attributed to the predominance of Mushroom Body neurons, which serve diverse roles in the larval phase of the fly’s ecology. Following metamorphosis, a more specialized core comprised of Antennal Lobe neurons emerges, reflecting the crucial role of olfaction in various aspects of adult fruit fly life. *D. melanogaster* undergoes complete metamorphosis, comprising four stages: egg, larva, pupa, and adult. The evolutionary theory posits that metamorphosis evolved to mitigate resource competition between young and adults, enabling larval and adult forms to occupy distinct ecological niches [51]. The significant distinction between the topology of the brain network of the two different phases of the *D. melanogaster* is in agreement with recent findings that some of the neuronal cells survive the metamorphosis but their connection completely changes and there is no evidence of the larval memory persisting into adulthood [12]. Our comprehensive comparison of adult and larval connectomes offers valuable insights into the intricacies of developing brain networks in *D. melanogaster*. It serves as a foundational framework for understanding the mechanisms underlying brain network development and provides a platform for future investigations.

## 4 Methods

### Network Properties

The brain networks are collected from [10, 52]. The nodes in these networks are the different neuronal cells and the edges are the synaptic connections between them. From both datasets, we take the table displaying the connections and constructed binary networks corresponding to the larval and adult networks using the Python package, networkx [53]. The adjacency matrix of each network is denoted as *A*, and the elements of this matrix *A*_*ij*_ = 1 if there is any synapse from neuron *i* to *j*. Both the data sets are directed and hence, *A*_*ij*_ ≠ *A*_*ji*_. The fundamental structural measure of a network is the node degree (*k*_*i*_ = ∑_*j*_ *A*_*ij*_), which quantifies the number of edges linked to a given node. The density measures how well connected the nodes are within the network and is defined as the ratio of the number of actual edges present in the network to the total number of possible edges. For a directed network, it is given as; 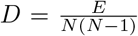, where *E* is the total number of edges and *N* is the total number of nodes present in the network. Further, the directed clustering for a node *i* is defined as [54]: 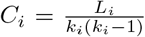, where, *L*_*i*_ is the number of directed links between the *k*_*i*_ neighbors of node *i, k*_*i*_ = *k*_*in*_ + *k*_*out*_ is the sum of the in- and out-degree of node *i*. All these structural properties were calculated with the help of the Python library networkx [53]. Further, the global efficiency is the average of the efficiencies of all the pairs of nodes in a network, given as follows; 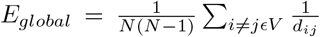. The Python package igraph is utilized for efficiency calculations. Further, a good measure of the influence of a node over the entire network is its betweenness centrality. We use betweenness centrality to compare the impact of the core neurons between the larva and adult brain network. For a node *v*, it is defined as; 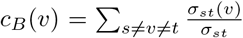, where, ∑*σ*_*st*_ is the number of the shortest path between all the pair of nodes *s* and *t* and *σ*_*st*_(*v*) is number of shortest paths that pass through the node *v*.

Further, we study community detection using the Louvain algorithm. In the case of larval connectome, 100 instances of community detection are carried out using 100 random seed values at the default resolution. Then we count the number of times any two nodes feature in the same community and make a matrix of it. Lastly, we run a community detection algorithm on this consensus adjacency matrix to obtain the final list of communities. In the case of the adult brain network, the computational complexity arising from the vast size of the resulting consensus matrix and the iterative process of updating the matrix with co-occurrences for each pair of nodes rendered the execution of a consensus community detection infeasible. Consequently, we opted to employ a single instance of community detection at the default resolution. The partitions obtained from the community detection of both brain networks are then used to obtain the value of modularity. Modularity for a directed graph is defined as [55]: 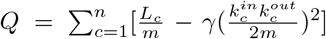, where, the sum is taken over all the *n* communities, *L*_c_ is the number of intra-community links for the community *c, m* is the total number of edges in the network, *γ* is the resolution parameter and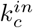 and 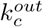 are the sum of in-degrees and out-degrees respectively in community *c*.

### Degree distribution and fitting

For fitting the degree distributions to the Weibull and power-law model, we use the Python power-law package created by Alstott et al. [56] which utilizes the statistical techniques laid down by Clauset et al. [31]. Simply put, the optimum minimum value of data (*x*_*min*_) is selected for which the Kolmogorov-Smirnov distance is the least. Then, the value of the exponent is calculated using Maximum Likelihood Estimation (MLE). MLE gives a more accurate estimation of the power law exponent than linear regression on doubly logarithmic axes because the latter has issues with hard-to-estimate errors, unreliable statistical measures (*r*^2^ values), and lack of normalization [31].

### Dcore Decomposition method

The Dcore decomposition is carried out using the algorithm laid out in the paper by Giatsidis *et al*.[27], applied to both the larval and adult connectomes. Given a graph *G*, we can obtain a unique D-core(*k, l*) (*DC*_*k,l*_) which is a maximal sub-graph that contains nodes of in-degree ≥ *k* and out-degree ≥ *l*. D-core is obtained for all pairs of *k* and *l* values such that for each pair, k ≤ *k*_*max*_ + 1 and l ≤ *l*_*max*_ + 1. Subsequently, we define D-core matrix of our network *G* as, *A*_*G*_(*k, l*) = (*dc*_*k,l*_)_*k,lϵN*_ where *dc*_*k,l*_ is the size of the D-core(k, l) (see Figure 2 (a)). We implement the algorithm mentioned in the supplementary material for all pairs of *k* and *l* values such that for each pair, k ≤ *k*_*max*_ + 1 and l ≤ *l*_*max*_ + 1. After obtaining the D-cores for all the pairs of k and l values, we define the D-core matrix of our network *G* as, *A*_*G*_(*k, l*) = (*dc*_*k,l*_)_*k,lϵN*_ where *dc*_*k,l*_ is the size of the D-core(k, l). The D-core matrix is drawn till *l*_*max*_ + 1 and *k*_*max*_ + 1 on the x and y axis respectively where *l*_*max*_ is the value of *l* such that *DC*_0,*lmax*_ is the last non-empty graph along the x-axis and similarly, *k*_*max*_ is the value of *k* such that *DC*_*kmax*,0_ is the last non-empty graph along the y-axis (see Figure 2 (a)). The last non-empty D-core, w.r.t. increase in *k* and *l* increase are highlighted with white colored oblique lines in the Figure 2 (b), (c)). These are termed as the frontier D-cores.

### Rich-club analysis

We calculate the in-degree and out-degree rich-club coefficient (RCC) defined as follows [24]:

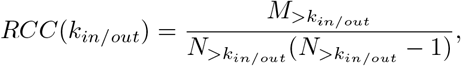

where, 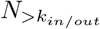 is the number of nodes with in/out-degree *> k*_*in/out*_ and 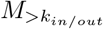 is the number of edges between them. As the high-degree nodes naturally have more probability of connecting with nodes of any type of degree including the high-degree one, the normalization of the above to the corresponding degree sequence preserved randomized networks gives a better idea of the rich-clubs [57]. Therefore, we calculated the normalized RCC.

### Degree preserving random networks

To investigate if a graph truly shows rich club organization we normalize its rich club coefficient with that of degree-preserved random graphs. Such a mode of randomization only changes the arrangement of the edges keeping the degrees of the nodes unchanged. This is carried out by implementing the directed edge swap method in the networkx [53] package. For both networks, 100 degree preserved random networks are created.

### Targeted attack and random attacks

Attack experiments are carried out to discern the effect of the loss of core neurons in both brain networks. Targeted attacks involve the removal of the entire set of core neurons at the same time and then measuring the parameters. While for random attack, a set of randomly chosen nodes are removed spontaneously. The size of the randomly selected set of nodes is set as the size of the cores. For calculating the directed global efficiency, we consider the largest and strongest component, whereas for the average directed clustering coefficient, we consider the entire graph.

### Diversity Indices

To assess the diversity in the neuron types of second-order neighbors of the core neurons, we implement two diversity indices: Simpson’s index 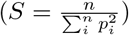 and Shannon index/entropy 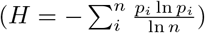. Where *p*_*i*_ = *N*_*i*_*/N* refers to the probability of a cell type, *n* is the total number of the cell types studied, *N*_*i*_ number of *i*^*th*^ cell type and *N* is the total number of the neurons studied. These measures were normalized to account for the different sizes. In the case of the Shannon index, this is done by taking the logarithm of the set of neurons being considered. The Shannon entropy is further utilized in assessing how the diversity of D-cores changed during the process of decomposition.

## 5 Acknowledgment

AS acknowledges the Department of Science and Technology (DST), Government of India, for financial support through grants DST/INSPIRE/04/2021/001893. PY acknowledges UGC for providing financial support under the Junior Research Fellowship scheme, as well as the lab members for their timely assistance and valuable discussions. We also express gratitude to IISER Tirupati for providing the High-Performance Computing facility. Additionally, we are thankful to Prof. Sitabhra Sinha for his insightful comments on this work.

## 1 Supplementary Material

### Degree Distribution

Here we show that the both in and out-degree of the larval well fit poorly with the power law and both the in and the out-degree of the adult brain fit poorly with the Weibull distribution (Fig. 1 (a), (b)). Further, the statistics of the maximal likelihood test performed to fit the data with the power-law distribution are tabulated in Table 1:

**Figure 1:**
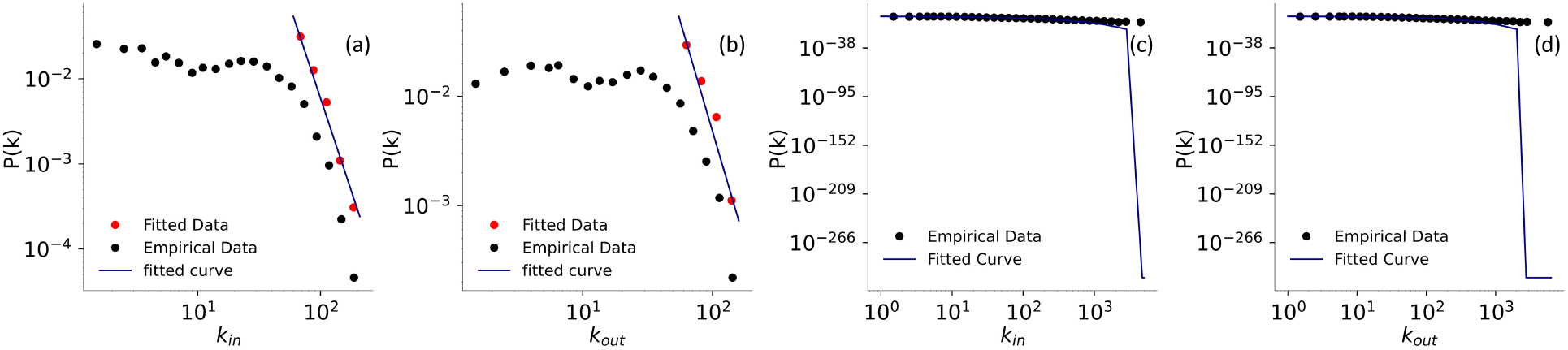
In and out-degree distribution of the larval (a, b) and the adult (c, d) brain networks. Here, showing the poor fitting of the larval in - and out-degree with the power-law and adult brain’s in-and out-degree with the Weibull distribution.

**Table 1:** Tests of Power Law behavior of degree distributions in Adult and Larva Connectome. LR stands for loglikelihood ratio. Positive values for the loglikelihood ratio (LR) indicate a preference for the power law model over alternative models when the p-value is less than 0.05. However, if the p-value exceeds 0.05, the sign of LR becomes an unreliable indicator of which model provides the better fit to the data.

|  |  | Adult |  | Larva |  |
| --- | --- | --- | --- | --- | --- |
|  |  | In-degree | Out-degree | In-degree | Out-degree |
| $\gamma$ | | 3.15 | 2.96 | 4.34 | 4.06 |
| **KS distance** |  | 0.01 | 0.02 | 0.05 | 0.08 |
| $x_{min}$ | | 113 | 90 | 60 | 55 |
| **Exponential** | LR | 8.09 | 6.33 | -3.30 | -8.38 |
| | $p$ | $6 \times 10^{-6}$ | $2.51 \times 10^{-10}$ | $9.72 \times 10^{-4}$ | $5.13 \times 10^{-17}$ |
| **Truncated Power Law** | LR | 0.19 | -0.17 | -4.14 | -9.52 |
| | $p$ | 0.97 | 0.82 | $5.02 \times 10^{-8}$ | $1.78 \times 10^{-8}$ |
| **Log normal** | LR | 1.73 | 0.0049 | -3.29 | -5.59 |
| | $p$ | 0.084 | 0.996 | 0.0009896 | $2.26 \times 10^{-8}$ |
| **Stretched Exponential** | LR | 3.8 | 2.0638 | -3.39 | -5.84 |
| | $p$ | $1.44 \times 10^{-4}$ | 0.0390 | 0.000696 | $5.29 \times 10^{-9}$ |
| **Lognormal Positive** | LR | 3.59 | 2.0965 | -3.29 | -5.59 |
| | $p$ | $3.27 \times 10^{-4}$ | 0.0360 | 0.0009896 | $2.26 \times 10^{-8}$ |

### Algorithm for the D-core decomposition

Dcore decomposition was carried out using the algorithm laid out in the paper by Giatsidis *et al*. [27]. Given a network called *G*, we can obtain a unique D-core(k, l), *DC*_*k,l*_ which is a maximal subgraph that contains nodes of in-degree ≥ k and out-degree ≥l. We implemented the algorithm in Python as follows:

INPUT: a directed graph G and positive integers k, l

OUTPUT: adjacency matrix corresponding to *DC*_*k,l*_, nodes belonging to *DC*_*k,l*_

1.1 initialise nodesarray

1.2 initialise adjacency matrix

1.3 recursion = True

2 while recursion == True and matrix is not empty

2.1 nodes to be deleted = nodes with either in-degree ≤ k OR out-degree l

2.2 remove the nodes to be deleted from the adjacency matrix

2.3 if nodes to be deleted is empty

2.3.1 recursion = False

end

end;

Further, we calculated the normalized Shannon entropy for all the Dcores of the Larval and the Adult brain network. The classification of neurons in the adult fly dataset is done using hierarchical annotations while the data for the larval connectome provides primarily the cell-type classification. Therefore, to have more overlap between the cell types of the larval connectome with the adul connectome, we have considered the ‘class’ annotation of the adult fruitfly dataset We plotted it in fig. 2. For both brains, the heterogeneity in the cell type increases as we move from the outer to the inner cells and then decreases. This decrement towards the inner cores is more in the larval brain as compared to the adult brain.

**Figure 2:**
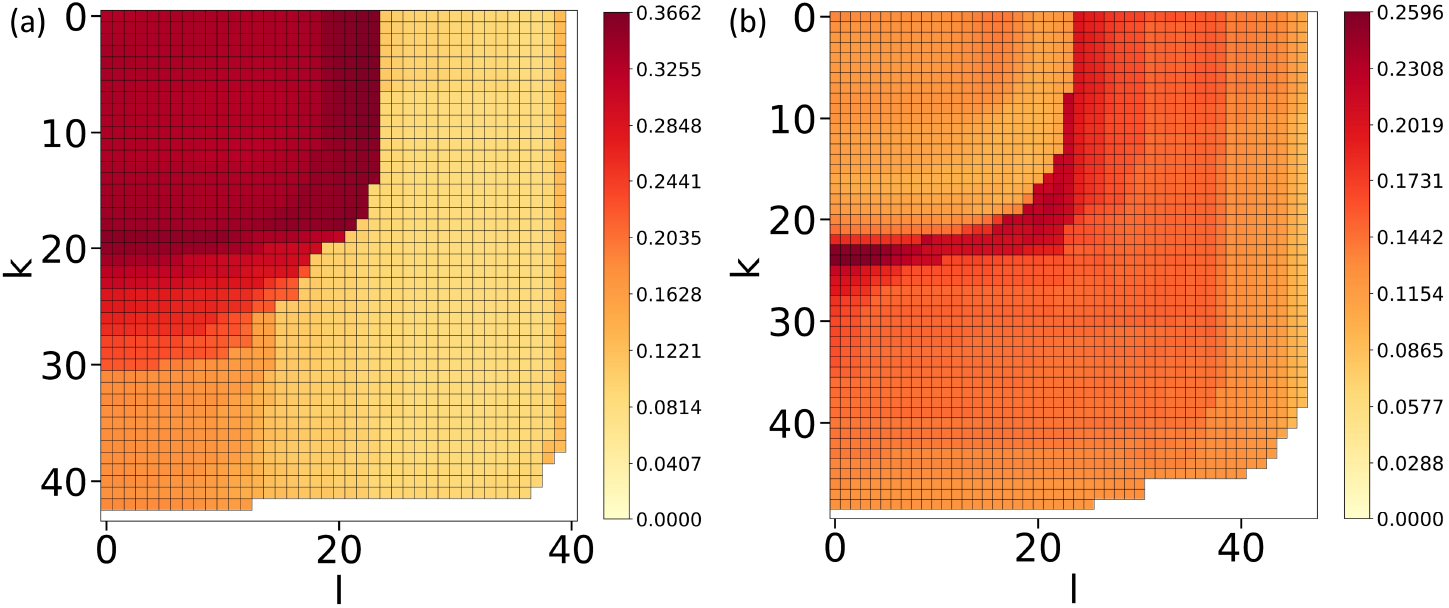
Normalized Shannon entropy for the different Dcore cells of the Larval (a) and the Adult (b) brain networks.

## References

[1] Olaf Sporns, Giulio Tononi, and Rolf Kötter. The human connectome: a structural description of the human brain. PLoS computational biology, 1(4):e42, 2005.

[2] Timothy EJ Behrens and Olaf Sporns. Human connectomics. Current opinion in neurobiology, 22(1): 144–153, 2012.

[3] Marryn M. Bennett Farzaan Salman Jacob D. Ralston Kaleb Hatch Raven F. Allen Alec M. Phelps Andrew P. Cook Jasper S. Phelps Mert Erginkaya Wei-Chung A. Lee Gwyneth M. Card Kevin C. Daly Andrew M. Dacks Han S.J. Cheong, Kaitlyn N. Boone. Organization of an ascending circuit that conveys flight motor state in drosophila. Current Biology, 34(5):1059–1075.e5, 2024. ISSN 0960-9822. doi: 10.1016/j.cub.2024.01.071.

[4] Alex Fornito, Andrew Zalesky, and Edward Bullmore. Fundamentals of brain network analysis. Academic press, 2016.

[5] Pew-Thian Yap, Guorong Wu, and Dinggang Shen. Human brain connectomics: networks, techniques, and applications [life sciences]. IEEE Signal Processing Magazine, 27(4):131–134, 2010.

[6] Barbara H Jennings. Drosophila–a versatile model in biology & medicine. Materials today, 14(5): 190–195, 2011.

[7] Marla B Sokolowski. Drosophila: genetics meets behaviour. Nature Reviews Genetics, 2(11):879–890, 2001.

[8] Lyubov Veverytsa and Douglas W Allan. Temporally tuned neuronal differentiation supports the functional remodeling of a neuronal network in drosophila. Proceedings of the National Academy of Sciences, 109(13):E748–E756, 2012.

[9] Shiri P Yaniv and Oren Schuldiner. A fly’s view of neuronal remodeling. Wiley Interdisciplinary Reviews: Developmental Biology, 5(5):618–635, 2016.

[10] Michael Winding, Benjamin D Pedigo, Christopher L Barnes, Heather G Patsolic, Youngser Park, Tom Kazimiers, Akira Fushiki, Ingrid V Andrade, Avinash Khandelwal, Javier Valdes-Aleman, et al. The connectome of an insect brain. Science, 379(6636):eadd9330, 2023.

[11] Dorkenwald S Matsliah A Sterling AR Schlegel P Yu SC McKellar CE Costa M Eichler K Bates AS Eckstein N Funke J Jefferis GSXE Murthy M. Lin A, Yang R. Network statistics of the whole-brain connectome of drosophila. bioRxiv [Preprint]. Feb 28:2023.07.29.551086., 2024. doi: 10.1101/2023.07.29.551086.

[12] James W Truman, Jacquelyn Price, Rosa L Miyares, and Tzumin Lee. Metamorphosis of memory circuits in drosophila reveals a strategy for evolving a larval brain. Elife, 12:e80594, 2023.

[13] Doe CQ. Lee K. A locomotor neural circuit persists and functions similarly in larvae and adult drosophila. Elife, 2021. doi: 10.7554/eLife.69767.

[14] Mark Newman. Networks. Oxford university press, 2018.

[15] Stephen B Seidman. Network structure and minimum degree. Social networks, 5(3):269–287, 1983.

[16] Seung Wook Oh, Julie A Harris, Lydia Ng, Brent Winslow, Nicholas Cain, Stefan Mihalas, Quanxin Wang, Chris Lau, Leonard Kuan, Alex M Henry, et al. A mesoscale connectome of the mouse brain. Nature, 508(7495):207–214, 2014.

[17] Fragkiskos D Malliaros, Christos Giatsidis, Apostolos N Papadopoulos, and Michalis Vazirgiannis. The core decomposition of networks: Theory, algorithms and applications. The VLDB Journal, 29:61–92, 2020.

[18] Nivedita Chatterjee and Sitabhra Sinha. Understanding the mind of a worm: hierarchical network structure underlying nervous system function in c. elegans. Progress in brain research, 168:145–153, 2007.

[19] Francesca Arese Lucini, Gino Del Ferraro, Mariano Sigman, and Hernán A Makse. How the brain transitions from conscious to subliminal perception. Neuroscience, 411:280–290, 2019.

[20] Emma K Towlson, Petra E Vértes, Sebastian E Ahnert, William R Schafer, and Edward T Bullmore. The rich club of the c. elegans neuronal connectome. Journal of Neuroscience, 33(15):6380–6387, 2013.

[21] Marcel A de Reus and Martijn P van den Heuvel. Rich club organization and intermodule communication in the cat connectome. Journal of Neuroscience, 33(32):12929–12939, 2013.

[22] Gorka Zamora-López, Changsong Zhou, and Jürgen Kurths. Cortical hubs form a module for multi-sensory integration on top of the hierarchy of cortical networks. Frontiers in neuroinformatics, 4:613, 2010.

[23] Martijn P Van Den Heuvel and Olaf Sporns. Rich-club organization of the human connectome. Journal of Neuroscience, 31(44):15775–15786, 2011.

[24] Martijn P Van Den Heuvel, Olaf Sporns, Guusje Collin, Thomas Scheewe, René CW Mandl, Wiepke Cahn, Joaqúin Goñi, Hilleke E Hulshoff Pol, and René S Kahn. Abnormal rich club organization and functional brain dynamics in schizophrenia. JAMA psychiatry, 70(8):783–792, 2013.

[25] Guusje Collin, René S Kahn, Marcel A De Reus, Wiepke Cahn, and Martijn P Van Den Heuvel. Impaired rich club connectivity in unaffected siblings of schizophrenia patients. Schizophrenia bulletin, 40(2):438–448, 2014.

[26] Randy L Buckner, Jorge Sepulcre, Tanveer Talukdar, Fenna M Krienen, Hesheng Liu, Trey Hedden, Jessica R Andrews-Hanna, Reisa A Sperling, and Keith A Johnson. Cortical hubs revealed by intrinsic functional connectivity: mapping, assessment of stability, and relation to alzheimer’s disease. Journal of neuroscience, 29(6):1860–1873, 2009.

[27] Christos Giatsidis, Dimitrios M Thilikos, and Michalis Vazirgiannis. D-cores: measuring collaboration of directed graphs based on degeneracy. Knowledge and information systems, 35(2):311–343, 2013.

[28] Sven Dorkenwald, Arie Matsliah, Amy R Sterling, Philipp Schlegel, Szi-Chieh Yu, Claire E McKellar, Albert Lin, Marta Costa, Katharina Eichler, Yijie Yin, et al. Neuronal wiring diagram of an adult brain. bioRxiv, 2023.

[29] Philipp Schlegel, Yijie Yin, Alexander S Bates, Sven Dorkenwald, Katharina Eichler, Paul Brooks, Daniel S Han, Marina Gkantia, Marcia Dos Santos, Eva J Munnelly, et al. Whole-brain annotation and multi-connectome cell typing quantifies circuit stereotypy in drosophila. bioRxiv, pages 2023–06, 2023.

[30] Mark EJ Newman. Power laws, pareto distributions and zipf’s law. Contemporary physics, 46(5): 323–351, 2005.

[31] Aaron Clauset, Cosma Rohilla Shalizi, and Mark EJ Newman. Power-law distributions in empirical data. SIAM review, 51(4):661–703, 2009.

[32] Yifei Zhang, Xiaodan Chen, Xinyuan Liang, Zhijiang Wang, Teng Xie, Xiao Wang, Yuhu Shi, Weiming Zeng, and Huali Wang. Altered weibull degree distribution in resting-state functional brain networks is associated with cognitive decline in mild cognitive impairment. Frontiers in Aging Neuroscience, 12, 2021. ISSN 1663-4365. doi: 10.3389/fnagi.2020.599112. URL https://www.frontiersin.org/articles/10.3389/fnagi.2020.599112.

[33] Rohwedder A. Schleyer M. et al. Saumweber, T. Functional architecture of reward learning in mushroom body extrinsic neurons of larval drosophila. Nat Commun., 9:1104, 2018. doi: 10.1038/s41467-018-03130-1.

[34] Dennis Pauls, Mareike Selcho, Nanae Gendre, Reinhard F. Stocker, and Andreas S. Thum. Drosophila larvae establish appetitive olfactory memories via mushroom body neurons of embryonic origin. Journal of Neuroscience, 30(32):10655–10666, 2010. doi: 10.1523/JNEUROSCI.1281-10.2010. URL https://www.jneurosci.org/content/30/32/10655.

[35] Leslie B Vosshall and Reinhard F Stocker. Molecular architecture of smell and taste in drosophila. Annu. Rev. Neurosci., 30:505–533, 2007.

[36] Andreas S Thum and Bertram Gerber. Connectomics and function of a memory network: the mushroom body of larval drosophila. Current opinion in neurobiology, 54:146–154, 2019.

[37] Katharina Eichler, Feng Li, Ashok Litwin-Kumar, Youngser Park, Ingrid Andrade, Casey M Schneider-Mizell, Timo Saumweber, Annina Huser, Claire Eschbach, Bertram Gerber, et al. The complete connectome of a learning and memory centre in an insect brain. Nature, 548(7666):175–182, 2017.

[38] Louis K Scheffer, C Shan Xu, Michal Januszewski, Zhiyuan Lu, Shin-ya Takemura, Kenneth J Hayworth, Gary B Huang, Kazunori Shinomiya, Jeremy Maitlin-Shepard, Stuart Berg, et al. A connectome and analysis of the adult drosophila central brain. Elife, 9:e57443, 2020.

[39] Nobuaki K Tanaka, Keita Endo, and Kei Ito. Organization of antennal lobe-associated neurons in adult drosophila melanogaster brain. Journal of Comparative Neurology, 520(18):4067–4130, 2012.

[40] Rachel I Wilson. Early olfactory processing in drosophila: mechanisms and principles. Annual review of neuroscience, 36:217–241, 2013.

[41] Philipp Schlegel, Alexander Shakeel Bates, Tomke Stürner, Sridhar R Jagannathan, Nikolas Drummond, Joseph Hsu, Laia Serratosa Capdevila, Alexandre Javier, Elizabeth C Marin, Asa Barth-Maron, et al. Information flow, cell types and stereotypy in a full olfactory connectome. Elife, 10:e66018, 2021.

[42] Hui-Hao Lin, Jason Sih-Yu Lai, An-Lun Chin, Yung-Chang Chen, and Ann-Shyn Chiang. A map of olfactory representation in the drosophila mushroom body. Cell, 128(6):1205–1217, 2007.

[43] Feng Li, Jack W Lindsey, Elizabeth C Marin, Nils Otto, Marisa Dreher, Georgia Dempsey, Ildiko Stark, Alexander S Bates, Markus William Pleijzier, Philipp Schlegel, et al. The connectome of the adult drosophila mushroom body provides insights into function. Elife, 9:e62576, 2020.

[44] Lalanti Venkatasubramanian and Richard S Mann. The development and assembly of the drosophila adult ventral nerve cord. Current opinion in neurobiology, 56:135–143, 2019.

[45] Claire Eschbach, Akira Fushiki, Michael Winding, Casey M Schneider-Mizell, Mei Shao, Rebecca Arruda, Katharina Eichler, Javier Valdes-Aleman, Tomoko Ohyama, Andreas S Thum, et al. Recurrent architecture for adaptive regulation of learning in the insect brain. Nature Neuroscience, 23(4):544–555, 2020.

[46] Claire Eschbach, Akira Fushiki, Michael Winding, Bruno Afonso, Ingrid V Andrade, Benjamin T Cocanougher, Katharina Eichler, Ruben Gepner, Guangwei Si, Javier Valdes-Aleman, et al. Circuits for integrating learned and innate valences in the insect brain. Elife, 10:e62567, 2021.

[47] Stephan Gerhard, Ingrid Andrade, Richard D Fetter, Albert Cardona, and Casey M Schneider-Mizell. Conserved neural circuit structure across drosophila larval development revealed by comparative connectomics. Elife, 6:e29089, 2017.

[48] Nathan C Klapoetke, Aljoscha Nern, Edward M Rogers, Gerald M Rubin, Michael B Reiser, and Gwyneth M Card. A functionally ordered visual feature map in the drosophila brain. Neuron, 110(10): 1700–1711, 2022.

[49] Shi Z. Gong Z. He S. Zhao, H. Modeling the evolution of biological neural networks based on caenorhabditis elegans connectomes across development. Entropy (Basel, Switzerland), 25(1):51, 2022. doi: 10.3390/e25010051.

[50] Gerhard Martin Technau. Brain development in Drosophila melanogaster, volume 628. Springer Science & Business Media, 2009.

[51] Hanna Ten Brink, André M de Roos, and Ulf Dieckmann. The evolutionary ecology of metamorphosis. The American Naturalist, 193(5):E116–E131, 2019.

[52] Codex: Connectome data explorer.

[53] Aric Hagberg, Pieter Swart, and Daniel S Chult. Exploring network structure, dynamics, and function using networkx. Technical report, Los Alamos National Lab.(LANL), Los Alamos, NM (United States), 2008.

[54] Giorgio Fagiolo. Clustering in complex directed networks. Physical Review E, 76(2):026107, 2007.

[55] Aaron Clauset, Mark EJ Newman, and Cristopher Moore. Finding community structure in very large networks. Physical review E, 70(6):066111, 2004.

[56] Jeff Alstott, Ed Bullmore, and Dietmar Plenz. powerlaw: a python package for analysis of heavy-tailed distributions. PloS one, 9(1):e85777, 2014.

[57] Vittoria Colizza, Alessandro Flammini, M Angeles Serrano, and Alessandro Vespignani. Detecting rich-club ordering in complex networks. Nature physics, 2(2):110–115, 2006.

